# Explaining Deep Neural Networks for the Prediction of Translation Rates

**DOI:** 10.1101/2023.06.02.543405

**Authors:** Frederick Korbel, Ekaterina Eroshok, Uwe Ohler

## Abstract

A recent convolutional neural network model accurately quantifies the relationship between massively parallel synthetic 5’ untranslated regions (5’UTRs) and translation levels, but the underlying sequence determinants remain elusive. Applying model interpretation, we extract representations of regulatory logic, revealing a complex interplay of regulatory sequence elements. Guided by insights from model interpretation, we adapt the model by human reporter data to obtain superior performance, which will promote applications in synthetic biology and precision medicine.

---

Regulation of mRNA translation enables rapid and local control of gene expression. As rate-limiting step, translation initiation is primarily controlled by the 5’ untranslated region (5’UTR) [1], which has a median length of 218 nucleotides in humans [2]. In it, regulatory sequence elements including RNA structural motifs and upstream open reading frames (uORFs) dictate the efficiency of translation. uORFs are present in 50 percent of human mRNA transcripts and conserved across species [3]. High-throughput genomics protocols that quantify translation have shown that uORFs are pervasively translated in a contextdependent fashion [4, 5], and that they both can promote or repress translation of the main ORF [6, 7].

Massively parallel reporter assays (MPRA) can be applied to measure the effect of vast numbers of designed and natural regulatory sequences on gene expression [8, 9]. A recent study built an MPRA library of entirely random 5’UTR fragments (i.e. uniform probabilities for A/C/G/U), assayed their effect on translation via the readout of mean ribosome load (MRL) observed in a human sample cell line, and used the resulting data to train a convolutional neural network (CNN) [10]. While successfully capturing the relationship between the 50-nucleotide long synthetic reporter sequences and translation, the underlying biological features that the model utilized remain unknown.

Explainable artificial intelligence (XAI) is a rapidly expanding focus area that aims at understanding complex machine learning models such as CNNs. To uncover the input features most important for prediction, feature attribution methods compute importance scores for input features, thus allowing to explain prediction output with respect to its input [11]. Hence, model interpretation can be applied as a tool to uncover functional sequence patterns and generate novel biological hypotheses [12]. To extract representations of translation regulatory logic, we apply integrated gradients [13] both to the original CNN fitted on the synthetic data (the “synthetic model”, dubbed Optimus-5-Prime by [10]) as well as a version that we trained on a second MPRA data set from the same study, consisting of human 5’UTR fragments (the “human model”, Fig. 1A). Moreover, we compare the underlying features driving translation prediction on synthetic and human 5’UTR sequences and make use of this knowledge to learn a model with superior performance.

**Figure 1:**
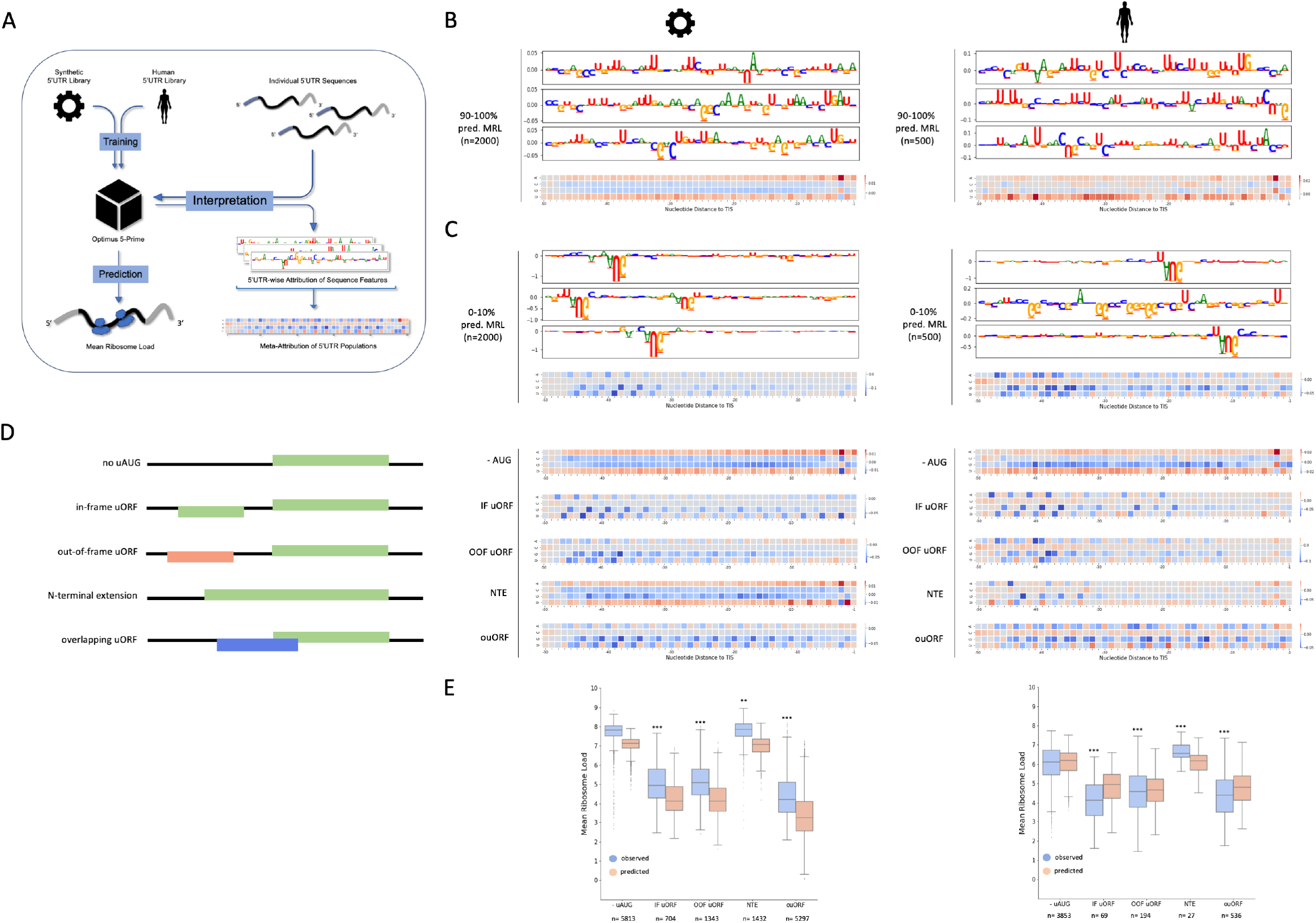
Extracting Learned Representations of Regulatory Logic. **(a)** A CNN predicts MRL of an mRNA given the sequence of its 5’UTR. We apply the integrated gradients algorithm on models trained on synthetic or human reporter data. From the explanations of predictions for individual 5’UTRs, we generate meta-attribution maps of 5’UTR populations. **(b)(c)** Feature attribution of the synthetic model trained and tested on synthetic reporters (left) and a human model trained and tested on human reporter data (right). Meta-attribution maps of the 10 percent of 5’UTRs with highest (b) or lowest (c) predicted MRL are complemented by example attribution maps of three individual 5’UTRs from the respective population. **(d)** Learned impact of putative translation in the 5’UTR on predicted MRL of the reporter mRNA for both models. Stratified by presence of one respective uORF or upstream start codon (uAUG) in the 5’UTR. **(e)** Observed vs. learned quantitative impact of upstream translation on MRL of the reporter mRNA for both models. Populations are the same as in D (unpaired two-tailed t-test; ** = p*<*0.01; ***= p*<*0.001).

We first computed meta-attribution maps by averaging feature importance values over a population of input samples to extract general features learned by the CNNs (Figure 1A). Isolating features that contribute positively or negatively to the prediction outcome, this approach readily verifies current knowledge. Both models learn that presence of the Kozak consensus at the start of the coding sequence (CDS) leads to increased ribosome load (Figure 1B) [14]. Moreover, high GC content in 5’UTRs is often associated with low levels of translation due to more stable secondary structures [2]. Both models capture this effect and positively attribute AU nucleotides while negatively attributing GC nucleotides (Figure 1B) throughout the sequence.

Upstream ORFs regulate translation of the main ORF depending on position and nucleotide context of their start and stop codons as well as their reading frame [7]. Both human and synthetic-data models reproduce these previously studied regulatory features, such as a global repressive effect of upstream translation, while strongly attributing the nucleotide context of initiation codons (Figure 1C). Furthermore, uORFs starting between -32 and -47 nucleotides upstream of the translation initiation site (TIS) were learned to be most repressive of ribosome load (Figure 1C,D), as long as their stop codon was also located upstream of the CDS. Translation of these uORFs may lead to a lower efficiency of ribosomal re-initiation or decay of the mRNA induced by ribosome stalling [15, 16]. In addition, a regular periodic positive attribution of A/U nucleotides suggests that translation from near-cognate start codons is learned by the neural networks (Figure 1B,D) [17].

To gain further insights into the effect of uORFs, we compared meta-attribution of reporters without upstream translation to reporters containing exactly one canonical upstream start codon (uAUG) or complete uORF (Figure 1D). The meta-attribution of the synthetic model exhibits strongly negative effects, while interpretation of the human MRL model identifies a low number of uORFs/uAUGs in the context of naturally occurring sequences. Consequently, the synthetic model overestimates the negative effect of upstream translation (Figure 1E). Nevertheless, both models distinguish between known regulatory mechanisms of upstream translation. For instance, in-frame uAUGs are not negatively attributed, since they can serve as an alternative TIS leading to an N-terminally extended protein product (NTE), which does not affect or may marginally increase ribosomal load of the coding sequence (Figure 1D,E). On the other hand, translation at uORFs and overlapping uORFs (ouORFs) is observed to negatively impact ribosome load of the CDS (Figure 1D,E), in line with current understanding [7].

Modeling mRNA translation with random synthetic 5’UTR sequences is attractive because of two reasons: First, it is unbiased, and in principle, the entire range of possible regulatory motifs is represented. Second, huge amounts of different random sequences can be generated. However, even an MPRA set of hundreds of thousands of sequences represents a tiny fraction of possible UTRs. When exploring the feature space in an entirely random fashion, various characteristics of endogenous, functional, evolved genetic sequences may therefore be over- or underrepresented. In turn, this may lead to a model learning strong weights for features that rarely occur in biological systems, while not capturing other motifs under evolutionary selection. This is important to note, since variation in nucleotide composition of the 5’UTR has a profound effect on translation efficiency [18]. For instance, human 5’UTRs are depleted of uORFs located directly upstream the canonical translation initiation site and have a higher overall content of guanine and cytosine nucleotides associated with stable 5’UTR secondary structure (Supplemental Figure 1) [3].

We therefore aimed to reveal additional features by interpreting models trained after exclusion of reporters carrying one or more uAUG in their 5’UTR. This resulted in a significant performance drop for the synthetic, but not for the human MRL model (Figure 2A). Additionally, almost no 5’UTR reporters with low to medium MRL were left in the synthetic data after removal of uAUG-reporters, suggesting that naturally occurring repressive motifs are underrepresented in synthetic 5’UTRs. This is supported by meta-attribution analysis, as a model trained on human 5’UTRs relies to a much greater extent on GC and U motifs, likely reflecting more stable secondary structure (Figure 2B,C). Taken together, a model trained on the available synthetic data alone, i.e. on entirely random data that is sizeable yet covers a small amount of possible sequences, does not sufficiently explain 5’UTR-mediated translation regulation in humans.

**Figure 2:**
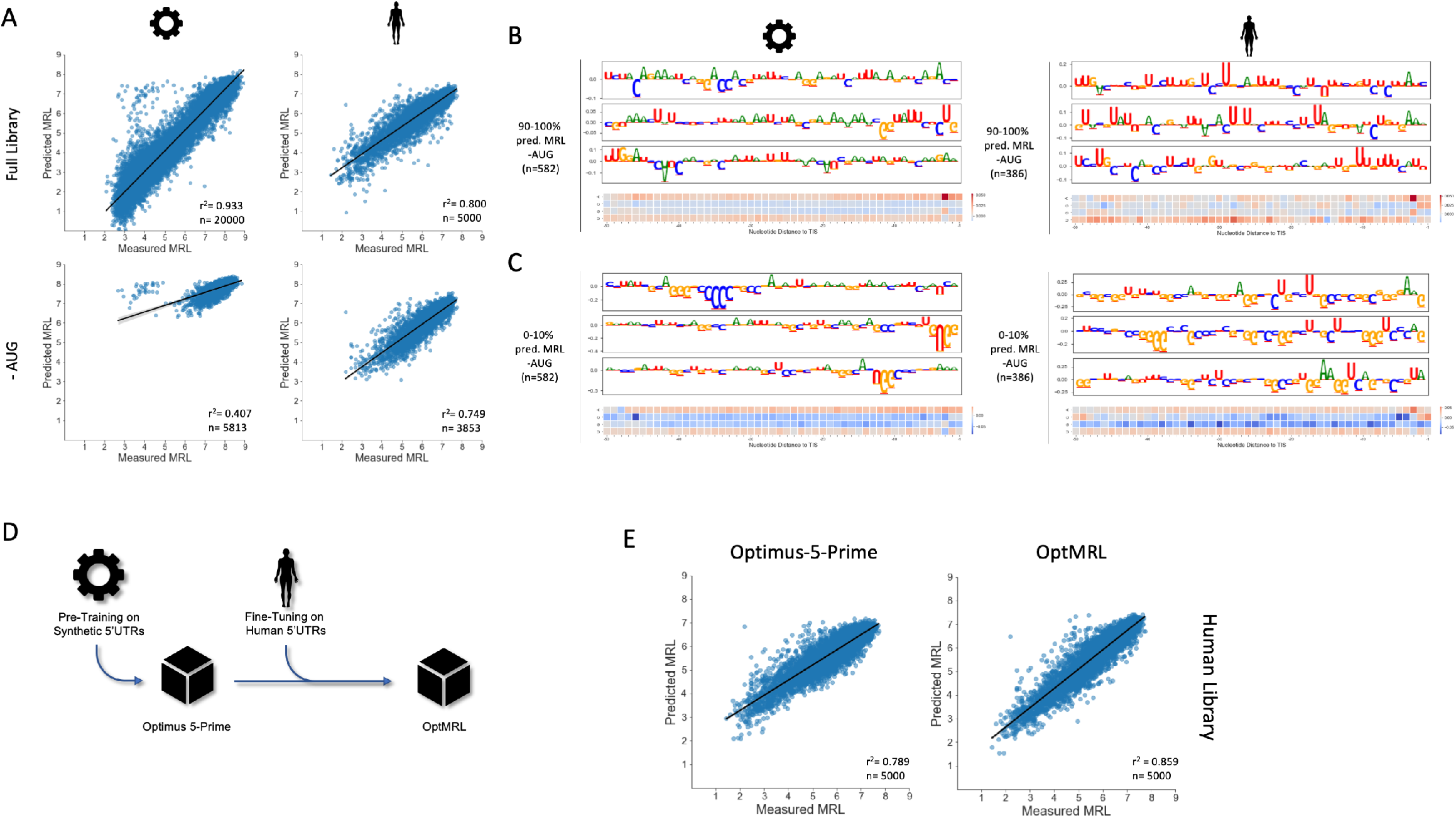
Developing an improved MRL Model with Interpretation and Transfer Learning. **(a)** Training and testing of the CNN on the synthetic (n=260,000) and human (n=20,000) reporter libraries and under exclusion of reporters with one or more AUG motif (-AUG). After removal of reporters with AUG in the 5’UTR, the performance of the synthetic model drops significantly, while it does not for the human model. **(b)(c)** Meta-attribution maps for CNNs trained and tested on synthetic (left) and human (right) data under exclusion of reporters with AUG in the 5’UTR. The 5’UTR reporters with highest 10 percent (b) and lowest 10 percent (c) predicted MRL are shown. **(d)** Retraining a model for prediction of ribosome load on human 5’UTRs. The best-performing model was chosen after additional training of the synthetic model on human data (n=20,000) for 1-15 epochs. **(e)** Performance of the synthetic model and an optimized MRL model (OptMRL) on a human 5’UTR reporter dataset (n=5,000). Additional training on human data reduced the prediction error on the human reporter library by one third.

To generate a model that benefits from the larger sample size of synthetic 5’UTR reporter sequences yet being sensitive to features present in human 5’UTRs, we re-trained the synthetic model on human data (Figure 2D). This transfer learning approach resulted in the OptMRL model, which more accurately reflects human 5’UTR sequence composition (Supplemental Figure 2) while conserving performance on synthetic 5’UTRs (Supplemental Figure 3). Importantly, it strongly improved on human data compared to the synthetic-only model, increasing prediction performance from 78,9 to 85,9 percent, thus reducing the error on the human 5’UTR reporter data by one third (Figure 2E).

The ability to extract representations of learned features from “black-box” models of gene regulation enables us to dissect model behavior, to understand characteristics of the available data (cf. Supplemental Figure 4), to put forth observations and hypotheses on gene-regulatory mechanisms, and to develop better models. Current developments in XAI aim to uncover interactions between individual features, and to shed light on higher order representations in the internal layers of the network. Yet, our work already demonstrates the impact of model interpretation for understanding and engineering RNA translation. We anticipate OptMRL to better capture endogenous translation regulation, which will promote our understanding of the effect of 5’UTR mutations and to engineer sequences with desired translational properties.

## Methods

### Overview of Models and Datasets

### Data Preprocessing

Datasets for computational analysis were downloaded from the Gene Expression Omnibus (GEO) under accession GSE114002 via the following link: https://www.ncbi.nlm.nih.gov/geo/query/acc.cgi?acc=GSE114002. Only reporters with a 5’UTR size of 50 nucleotides were considered in this study. Specifically, the unmodified eGFP library of synthetic 5’UTR reporters (GSM3130435) and the designed library containing human 5’UTR reporters (GSM3130443) were selected for analysis. Training and test datasets for the synthetic reporter library are the same as in Sample et al. [10] with a training set size of 260,000 and a test set size of 20,000. To assure direct comparability of the models, the preprocessing workflow of Sample et al. [10] was followed. Hence, the 5,000 human 5’UTR reporters with most sequencing reads across fractions were defined as test set and the remaining 20,000 human 5’UTR reporters were defined as training set.

### Model Training and Validation

All models investigated in this study follow the published Optimus-5-Prime architecture. They consist of three convolutional layers with 120 filters each, followed by a fully connected layer with 40 neurons and the output layer on top. If not stated otherwise, training hyperparameters were taken from Sample et al. [10]. Reporter-wise mean ribosome load values as target values for training of deep learning models have been adopted without change from GSE114002. As in Sample et al. [10], target values of training and test datasets were scaled separately using the scikit-learn [19] (v1.0.2) standardscaler before model fitting. Later, predicted target values were inverse-transformed accordingly. During training, loss on validation dataset was monitored after each epoch to review training history. The number of training epochs was determined manually based on the relationship between training and validation loss. All models were implemented in the tensorflow framework [20] (v2.4.3) with adopted code where indicated.

### Model Interpretation

Model interpretation was performed with the integrated gradients algorithm [13], where changes in prediction outcome are measured for small changes in the input. In a custom python script, the integrated gradients were computed over a linear path from a null matrix as baseline to the actual input matrix in 50 steps. The obtained values were then scaled relative to the input matrix. Attribution maps were obtained through visualization of attribution matrices as sequence logos, with letter height representing the relative importance of the individual nucleotide positions. Sequence logos were computed with code adapted from the concise package [21] and Ghanbari et al. [12].

To generate meta-attribution maps that summarize the feature-space of large populations of input examples, attribution matrices from individual input sequences were added and normalized by sample size. Given a number (n) of feature attribution matrices (N), the respective meta-attribution matrix (M) is obtained through:

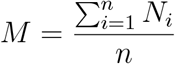

### Defined Subtypes of UTRs

To dissect the contribution of individual functional types of upstream translation on ribosome load, 5’UTR reporters were sorted into groups depending on their reading frame as well as the location of their downstream stop codon. We defined upstream open reading frames as a combination of an AUG codon and a downstream stop codon (UAA, UGA, UAG) within the same biological reading frame. A python script was used to sort and separate 5’UTR reporters into five groups. Reporters without uAUG in the 5’UTR were used as control group and those with more than one or ambiguous upstream translation signals were excluded from the analysis. Correspondingly, we distinguished between reporters with one non-overlapping uORF in-frame with the coding sequence (IF uORF), one non-overlapping uORF out-of-frame with the coding sequence (OOF uORF), one overlapping uORF (ouORF) as well as an N-terminal extension (NTE) of the protein product elicited by an in-frame uAUG.

### Occlusion of Sequence Features

In a two-step iterative process, subsets of 5’UTR reporters with certain features were removed from the training, testing and attribution workflow in order to reveal additional subtle sequence features. For both synthetic and human data, models were then fitted separately on the remaining data. In a first iteration, 5’UTR reporters with at least one uAUG were removed from the training and test datasets (-uAUG). After fitting a model on the remaining data, feature attribution was performed accordingly. As our XAI analysis led us to notice that the adapters used in [10] ended with an adenosine, reporters with ‘UG’ as the first two nucleotides in the 5’UTR were additionally removed from the datasets in the second iteration,(-uAUG -UG). Again, feature attribution was performed on the models fitted on the remaining data.

### Transfer Learning

The optimized MRL model was obtained through a transfer learning approach. To this end, the Optimus-5-Prime model published by Sample et al. [10], trained on synthetic 5’UTR reporters, was re-trained on 20,000 human 5’UTR reporters. To determine the optimal number of training epochs, a search was performed over a space of 1 to 15 training epochs using keras-tuner [22] (v1.1.3). The best-performing model was then selected for further analysis (OptMRL, Table 1).

**Table 1:** Overview of Model Training and Datasets

| Dataset | Training Size | Test Size | Model | Epochs |
| --- | --- | --- | --- | --- |
| synthetic (full) | 260,000 | 20,000 | O5P | 3 |
| synthetic (-uAUG) | 122,927 | 6,014 | MRL-uAUG | 3 |
| synthetic (-uAUG -UG) | 118,525 | 5,813 | MRL-uAUG-UG | 3 |
| human (full) | 20,000 | 5,000 | hMRL | 9 |
| human (-uAUG) | 14,564 | 3,942 | hMRL-uAUG | 7 |
| human (-uAUG -UG) | 14,079 | 3,853 | hMRL-uAUG-UG | 7 |
| synthetic + human (full) | 260,000 (s) + 20,000 (h) | 20,000 (s) + 5,000 (h) | OptMRL | 3 + 13 |

### Statistics

Outliers are dotted. P-values are the result of two-tailed t-tests calculated with scipy [23] (v1.8.0). Sample sizes and p-values are indicated where relevant. Statistical annotation (Supplemental Figure 1) was performed with the statannotations package [24] (v0.5.0). All boxplots display the median in their central part, interquartile range (25th/75th percentile) on the box edge, and 1,5x interquartile range as whiskers. The coefficient of determination, r-squared, was used as measure of performance throughout this study (e.g. how well are measured mean ribosome load values explained by the artificial neural networks).

## Code Availability

All code was developed in python (v3.9.10). Code and data generated in this study are deposited on github under the following link: https://www.github.com/ohlerlab/mlcis. The OptMRL model can also be downloaded there.

## Supplemental Information

**Supplemental Figure 1:**
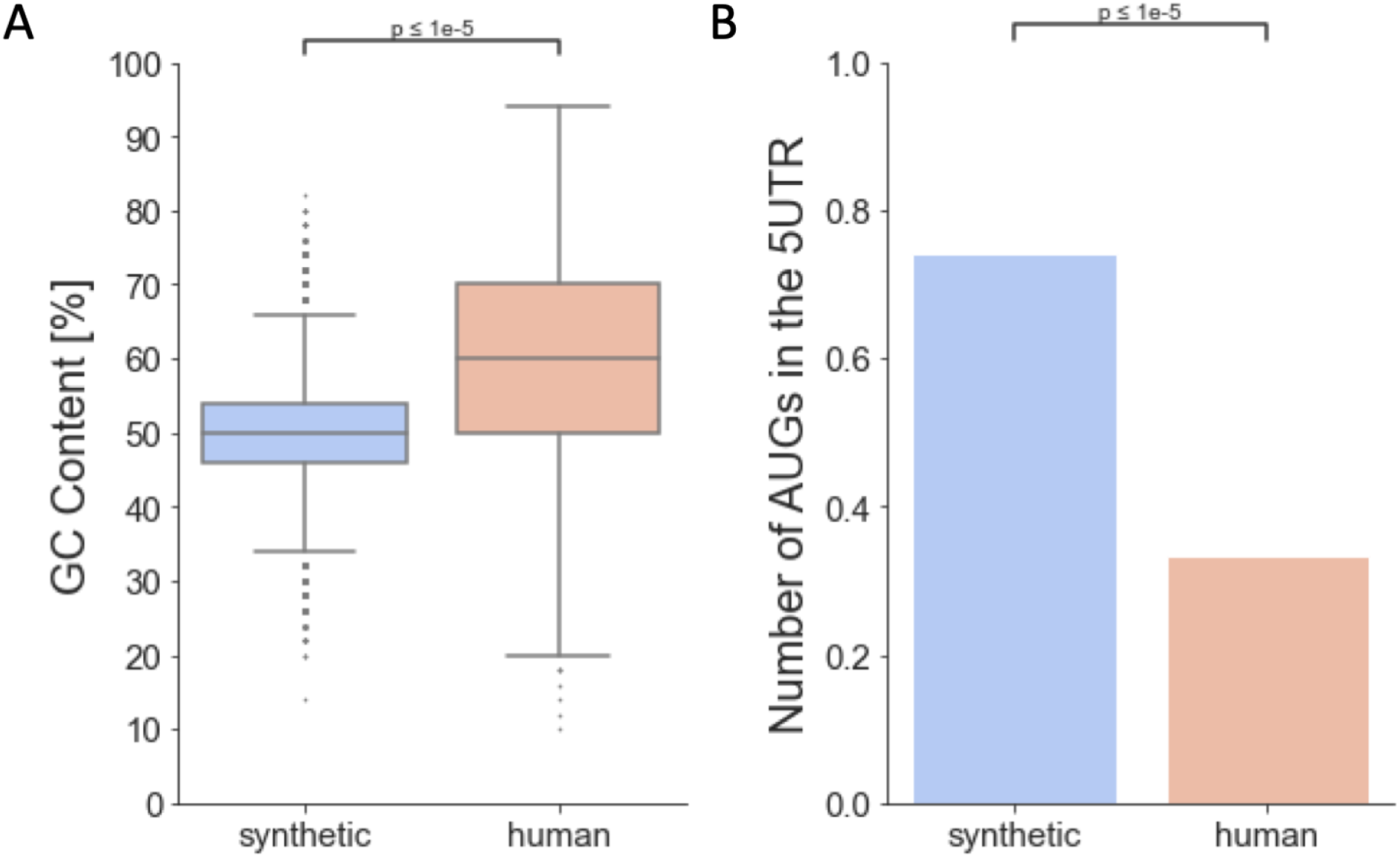
Sequence Composition of Synthetic and Human MPRA Libraries. Comparison between the synthetic 5’UTR library (n= 280,000) and the human 5’UTR library (n= 25,000) from Sample et al. [10]. **(a)** GC-content, which is indicative of secondary structure, is significantly higher for human 5’UTRs than for random synthetic 5’UTRs. **(b)** The fraction of 5’UTRs with uAUG is significantly lower for the human 5’UTR library, which are depleted for uAUGs right upstream the TIS, than for the synthetic 5’UTR library. Results of unpaired two-tailed t-tests are indicated.

**Supplemental Figure 2:**
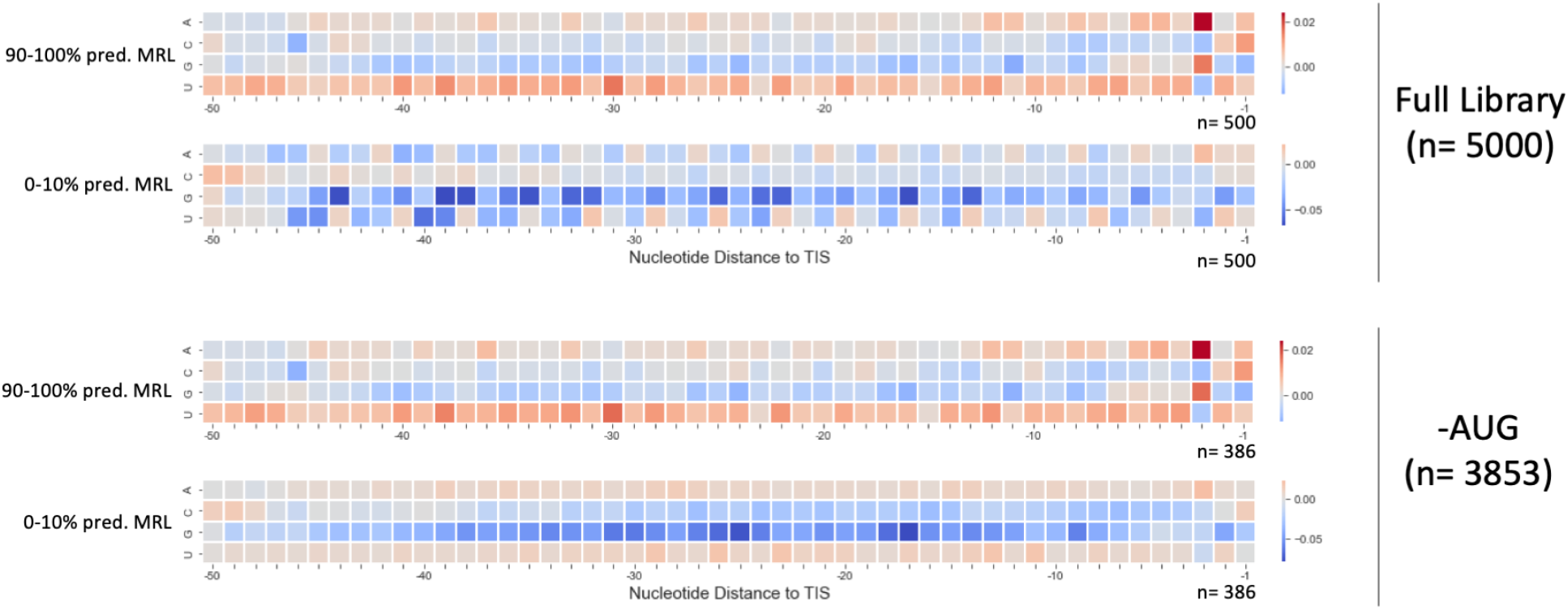
Meta-Attribution Maps of the OptMRL Model. Meta-attribution maps of the optimized MRL model for the full test library of human 5’UTRs and under exclusion of reporters with AUG in the 5’UTR. The model relies on general features of 5’UTRs when no AUG is present, predominantly on GC-rich patterns and U-rich patterns, i.e. likely related to secondary structure.

**Supplemental Figure 3:**
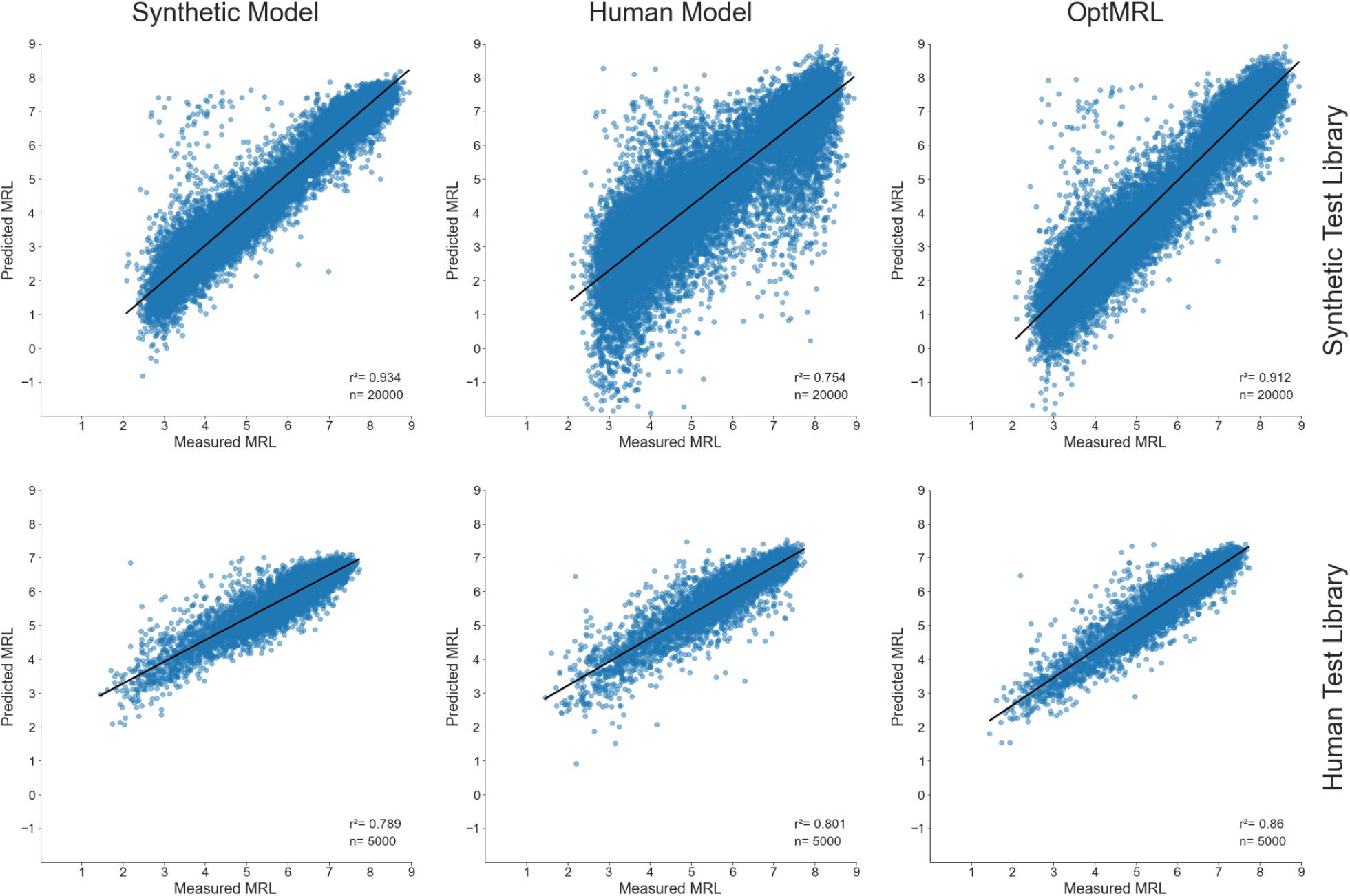
Performance of three CNNs on Synthetic and Human MPRA Libraries. The predictive performance of the synthetic model from Sample et al. [10], the human MRL model trained on 20,000 human 5’UTR reporters, and the optimized MRL model trained on 260,000 synthetic and re-trained on 20,000 human 5’UTR reporters, are compared on both a synthetic and a human test library. Although the human model performs better on a test library of human 5’UTR reporters than the synthetic model, performance is significantly lower on a synthetic 5’UTR test library. On the other hand, the optimized MRL model conserves performance on synthetic 5’UTR reporters, while increasing performance on human 5’UTR reporters compared to both other models.

**Supplemental Figure 4:**
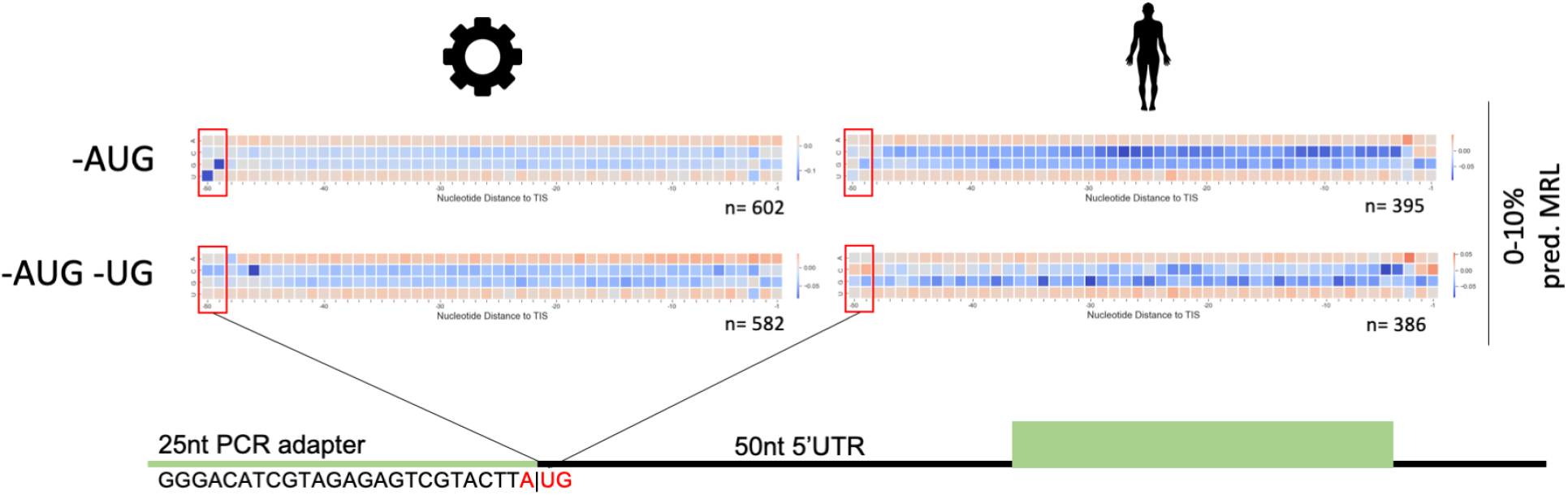
Meta-Attribution reveals an Experimental Artifact. Meta-attribution maps of the 10 percent of reporters with highest predicted ribosome load for versions of the model trained on the synthetic and human libraries, respectively. Models were trained and tested under exclusion of reporters with at least one upstream initiation codon (-AUG) and of those that exhibit ‘UG’ in the first two positions of their 5’UTR (-AUG -UG). A uridine and guanine as first two nucleotides in the 5’UTR of a reporter show strong attribution: together with the adenine in the last position of the PCR adapter, they create an AUG start codon in an alternative reading frame, resulting in a uORF overlapping into the coding sequence of those mRNAs. This artifact was learned by the model and is therefore revealed by feature attribution.

